# Glucose-regulated circular RNA Rabep1 regulates pancreatic β-cell growth by modulating miR-335-3p/PTEN axis

**DOI:** 10.1101/2024.06.24.600308

**Authors:** Debojyoti Das, Sharmishtha Shyamal, Smruti Sambhav Mishra, Susovan Sadhukhan, Amaresh C. Panda

**Affiliations:** Institute of Life Sciences, Nalco Square, Bhubaneswar, Odisha, India

**Keywords:** Pancreatic β-cells, diabetes, circular RNA, microRNA, insulin, glucose

## Abstract

**Highlights:**

- Identified circRNAs expressed in βTC6 cell line
- First report identifying glucose-regulated circRNAs in pancreatic β-cell
- *CircRabep1* regulates β-cell growth by binding to miR-335-3p

Circular RNAs (circRNAs) are a large family of closed-loop RNA molecules emerging as novel regulators of gene expression. Although several circRNAs are known to regulate various biological processes, the functions of most circRNAs expressed in pancreatic β-cells remain to be discovered. Since short-term glucose treatment induces pancreatic β-cell growth and promotes insulin production, we wanted to explore the role of glucose-regulated circRNAs in pancreatic β-cell physiology. Our RNA-seq analysis identified more than 300 differentially expressed circRNAs in high-glucose compared to low-glucose treated βTC6 cells. A subset of differentially expressed and abundant circRNAs was validated by various biochemical methods, including circular RNA *Rabep1* (*circRabep1*). Moreover, the downregulation of *circRabep1* in high glucose-treated βTC6 cells suggested a possible function in β-cell physiology. Furthermore, analysis of the *circRabep1*-miRNA-mRNA regulatory network discovered the association of *circRabep1* with miR-335-3p, a suppressor of *Pten* expression. Importantly, inhibition of miRNA function by miR-335-3p inhibitor results in upregulation of PTEN levels, suppressing β-cell growth and proliferation. Furthermore, silencing *circRabep1* decreased PTEN expression by sponging miR-335-3p, promoting cell proliferation. We propose that the downregulation of *circRabep1* in high-glucose treated β-cell leads to an increase in β-cell proliferation by suppressing PTEN expression through derepression of miR-335-3p.

Graphical Abstract
Schematic showing the molecular function of glucose-regulated *circRabep1* in pancreatic β-cell growth by binding to miR-335-3p.

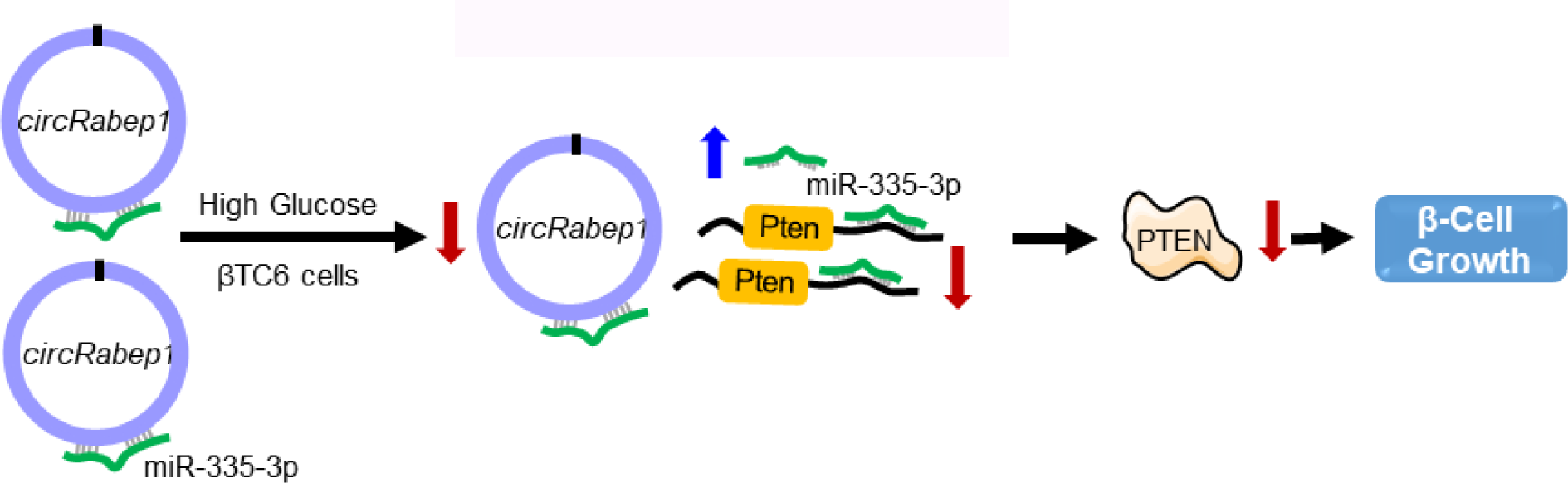

## INTRODUCTION

Diabetes mellitus (DM), characterized by defective glucose homeostasis, is a widespread epidemic in almost all countries ^1^. Diabetes is recognized as a predominant health threat leading to long-term complications in the kidney, heart, liver, eyes, and foot problems. Resistance to insulin by different organs or insufficient insulin production from pancreatic β-cells leads to diabetes ^2^. Numerous factors are responsible for the development of diabetes, including pancreatic β-cell death and impaired insulin production ^3–6^. In recent years, large-scale high-throughput sequencing identified numerous mutations and genes associated with the development of diabetes ^7^. Increasing evidence suggests that onset of diabetes affects insulin biosynthesis and β-cell death rate, which themselves are tightly regulated processes by transcription factors, RNA-binding proteins, miRNAs, long noncoding RNAs, and poorly characterized circular RNAs (circRNAs) ^6, 8–10^.

CircRNAs are a vast family of ubiquitously expressed covalently closed RNA molecules formed by the head-to-tail backsplicing of pre-mRNAs ^11, 12^. CircRNAs show tissue-specific expression, and many of them are evolutionarily conserved ^11, 13^. Due to the lack of free ends, circRNAs are very stable and resistant to endogenous RNA exonucleases. Although the molecular mechanism of gene regulation by most of the circRNAs is yet to be explored, four mechanisms well accepted till date include; acting as miRNA sponge, can act as a decoy for RBPs, can regulate transcription, and can be translated into proteins ^14^. Several circRNAs are known to regulate gene expression in normal physiology, and their deregulation leads to the development of diseases ^15^. Recent studies, including our previous study, highlighted the differential expression and association of circRNAs in pancreatic β-cells in response to a high-fat diet ^9,16^. Although the mechanism of several circRNAs in pancreatic β-cells has been characterized, we still have a limited understanding of the role of most of the circRNAs expressed in β-cells during the development of diabetes^9^.

Given that altered expression of circRNAs might be involved in β-cells gene regulation during the development of diabetes, we used high-throughput RNA sequencing to identify differentially expressed circRNAs in glucose-treated βTC6 cells. In this study, we performed RNA sequencing of βTC6 cells treated with high glucose compared to low glucose, wherein we identified a few abundant and differentially expressed circRNAs, including *circRabep1*. Furthermore, we characterized the *circRabep1*-miR-335-3p-PTEN regulatory network that regulates β-cells growth in high glucose conditions. Our findings thus help to unveil the molecular mechanism of circRNAs in β-cell physiology.

## MATERIALS AND METHODS

### Pancreatic islet isolation, βTC6 culture, and glucose treatment

Four to six-week-old male C57Bl/6 mice were acquired from the Institute of Life Sciences breeding colonies. The whole pancreas was dissected from the mice and washed in PBS, followed by islet preparation using collagenase digestion and the Ficoll gradient method, as described previously ^17^. The islets were cultured in RPMI media supplemented with 15 % FBS and antibiotics. The islets were cultured in Dulbecco’s modified eagle’s medium (DMEM) with 2.5 mM low glucose (LG) or 25 mM high glucose (HG) for 24 h, followed by RNA isolation. High glucose DMEM supplemented with 15 % FBS and antibiotics was used to culture the mouse βTC6 cells in 5% CO2 at 37°C. For the glucose treatment experiment, βTC6 cells were cultured in DMEM containing 5 mM glucose supplemented with 15 % FBS and antibiotics for a few weeks before glucose treatment. The βTC6 cells were treated with 25 mM high glucose (HG) or 2.5 mM low glucose (LG) DMEM for 24 h, followed by RNA isolation.

### Total RNA isolation, RT-PCR analysis, and DNA sequencing

Total RNA extraction from β-cells and pancreatic islets was performed using Trizol reagent. RNA concentration was quantified using Nanodrop 2000 or Multiskan sky spectrophotometer, followed by cDNA synthesis using Maxima RT (Thermo Fisher Scientific) or Luna Superscript master mix (NEB) following the manufacturer’s protocol. Specific circRNAs were PCR amplified with divergent primer pairs, as mentioned in **Supplementary Table S1** ^18^. DreamTaq Green PCR Master Mix was used to amplify the desired amplicon with a PCR condition of 95°C for 2 min, followed by 40 cycles of 95 °C for 5 s and 60 °C for 20 s. PCR products were resolved in 2% agarose gel previously stained with SYBR-Gold dye and visualized on an ultraviolet trans illuminator. The PCR products were purified, followed by Sanger sequencing using one of the divergent primers to confirm the back splicing junction (BSJ) sequence.

### Library preparation and RNA sequencing of glucose-treated βTC6 cells

The total RNA from the HG and LG treated βTC6 cells were rRNA depleted using NEBNext® rRNA Depletion Kit (Human/Mouse/Rat) (NEB), followed by fragmentation and the cDNA library preparation using the NEBNext® Ultra™ II Directional RNA Library Prep Kit (NEB) following the manufacturer’s protocol. The cDNA libraries were sequenced with single-end 150 bp reads on the Illumina NextSeq 550 platform using High Output Kit v2 (Illumina); data were deposited in ENA-Accession No. PRJEB64535.

FastQC was used to check the quality of the raw reads, followed by the removal of the adapters. The clean reads were aligned to the mouse genome (mm10) using the STAR aligner with default parameters. The mRNA expression in each sample was assessed by FeatureCounts (Subread package: v2.0.1). Differential gene expression was performed using edgeR ^19^. Low-expression transcripts were filtered out using edgeR filtering, and the heat map was generated using Z-score. For circRNA identification, the alignment information from STAR was used as an input to CIRCexplorer2 (v2.3.0) annotation module ^20^. The circRNA expression levels were calculated using transcripts per million (TPM; [circRNA read number/total reads in the sample x 1,000,000]). The differentially expressed circRNAs between HG and LG samples were identified using the t-test with a P-value ≤ 0.05. The complete list of circRNAs expressed in βTC6 cells and differential expression analysis can be found in **Supplementary Table S2.**

### RNase R treatment, Quantitative PCR, and Droplet Digital PCR (dd-PCR) analysis

To confirm the circularity of the selected circRNAs, total RNA from islets and βTC6 cells were treated with RNase R (Epicentre), followed by RT-qPCR analysis of linear and circRNAs as mentioned previously ^18^. PowerUp SYBR Green Master Mix (Thermo Fisher Scientific) was used to amplify specific circRNAs using its divergent primer pairs in QuantStudio 3/6 Real-Time PCR System (Thermo Fisher Scientific). The delta-CT method was used to check for the relative abundance of individual circular RNA molecules, and the data were normalized with *18s* rRNA or *Actb* mRNA. Digital droplet PCR (dd-PCR) was performed on total RNA isolated from βTC6 cells to quantify the absolute copy number of circular RNAs. One µg of total RNA was used to prepare cDNA using the Maxima reverse transcriptase. The dd-PCR reaction of 20 µl was designed to contain cDNA converted from 10 ng of RNA, 250 nM primers, and EvaGreen Supermix (Biorad) ^21^. The PCR droplets were generated with the QX200 droplet generator followed by PCR amplification at 95°C for 2 min and 45 cycles of 5 s at 95°C plus 60 s at 60°C followed by droplet stabilization for 5 min at 95 °C and 5 min at 4 °C. The PCR droplets were quantified with the QX200 droplet reader, and the number of circRNA per ng of RNA was calculated ^21^.

### RNA stability, transfection experiments, and WST-1 assay

The βTC6 cells were treated with 10 μg/mL actinomycin D (Sigma) for 0, 0.5, 1, 2, 4, 8, and 16 h, followed by total RNA isolation and RT-qPCR analysis to assess circRNA stability. To study the effect of *circRabep1* on *Pten* mRNA expression, the βTC6 cells were transfected twice at 24 h with *circRabep1* GapmeR or control GapmeR at 100 nM concentration using Lipofectamine RNAiMAX transfection reagent (Thermo Fisher Scientific). After two days, transfected βTC6 cells were treated with 10 μg/mL actinomycin D (Sigma) for 0h, 0.5h, 1 h, 2 h, 4 h, and 8 h, followed by RT-qPCR analysis of *Gapdh* and *Pten* mRNA. The *circRabep1* and PTEN expression were analyzed by qPCR or western blot analysis after 48 h of *circRabep1* GapmeR transfection. βTC6 cells were transfected with 100 nM of control or miR-335-3p inhibitor alone or along with *circRabep1* GapmeR for the rescue experiment. After 48 h of transfection, the media was changed, and 100 µL fresh culture media was added, followed by 10 µL of WST-1 reagent and incubated for 4 h at 37 °C, followed by measuring the absorbance at 450nm.

### CircRNA ASO Pulldown assay

The βTC6 cells at about 90% confluence were washed with PBS twice, followed by lysis with polysome extraction buffer (PEB) supplemented with RNase and protease inhibitors. The cell lysate was incubated for 90 min at 4°C with *circRabep1* backsplice junction-specific biotin-labeled ASO or control ASO, followed by incubation with streptavidin beads for 30 min at room temperature. The pulldown beads were washed thrice in ice-cold 1X TENT buffer, followed by RNA isolation and RT-qPCR analysis of *circRabep1* and miR-335-3p ^22^.

### Reporter assay

To confirm the regulation of PTEN by miR-335-3p, the 3’UTR of PTEN targeted by miR-335-3p was cloned downstream of renilla ORF in the psiCHECK2 vector. For reporter assay, βTC6 cells were transfected with control miRNA or miR-335-3p mimic along with psiCHECK2 and psiCHECK2-PTEN 3’UTR reporter plasmids. The renilla and firefly luciferase activities were measured using Dual-Glo Luciferase Assay System (Promega) after 16 h of transfection. The relative renilla luciferase activity was normalized to firefly luciferase activity.

### Western Blot

Two days after transfection with *circRabep1* GapmeR or miR-335-3p inhibitor, βTC6 cells were harvested and lysed with RIPA buffer supplemented with protease inhibitor. The lysate was resolved in a SDS-PAGE and transferred to a nitrocellulose membrane. The membrane was incubated with PTEN (Abcam) and GAPDH (CST) antibodies. After the overnight incubation, the blots were washed and incubated with appropriate HRP-conjugated secondary antibodies, followed by visualization using chemiluminescence reagents.

### Polysome analysis

Polysome profiling was performed on *circRabep1* knockdown cells to study *Pten* mRNA translation as described previously ^23^. Briefly, proliferating βTC6 cells transfected with control or *circRabep1* GapmeR were treated with cycloheximide (100µg/mL) for 10 min followed by washing with PBS and lysis using PEB supplemented with RNase inhibitor and protease inhibitor. The lysate was cleared by centrifugation, and the supernatant was loaded onto a 10-50% sucrose gradient followed by ultracentrifugation (Optima XPN-100, Beckman Coulter) at 1,90,000 x g (SW41Ti rotor) for 2.5 hours at 4°C. The sucrose gradient was collected into 12 fractions using the polysome fractionator (BioComp Gradient Station IP), followed by isolation of RNA and RT-qPCR analysis of target RNAs.

### Construction of the competing endogenous RNA (ceRNA) Network

To analyze the *circRabep1*-miRNA-mRNA regulatory networks, we predicted the putative miRNAs targeting *circRabep1* using miRDB and circAtlas database ^24, 25^. To construct the ceRNA network for the *circRabep1-*associated miRNAs, we identified the experimentally validated mRNA targets using the miRTarBase release 9.0 ^26^. The target site of miR-335-3p on *Pten* 3’UTR was predicted by miRTarBase using miRanda, TargetScan, and miRDB ^24, 26, 27^. The *circRabep1*-miRNA interactions and miRNA-mRNAs information was used to construct the ceRNA network of the *circRabep1*-miRNA-mRNA axis.

### Gene Ontology (GO) and Kyoto Encyclopedia of Genes and Genomes (KEGG) Pathway Analysis

The gene set enrichment and KEGG pathway analysis for the target genes in the *circRabep1*-miRNA axis were performed using KOBAS ^28–30^. The enrichment score for the GO terms and KEGG pathways were calculated as - LOG10 of the False Discovery Rate (FDR), and the values were plotted as bubble plots along with the gene ratio ^29,30^. In addition, the protein-protein interaction (PPI) network of the genes in the *circRabep1*-miRNA-mRNA axis was analyzed using the Cytoscape StringApp (3.10.0).

### Visualization and statistical analysis

GraphPad Prism 6.0 software or Microsoft Excel was used to plot the graphs for data visualization. All experiments were repeated at least 3 times and represented as the mean ± standard error of the mean (SEM). Statistical significance was calculated by student’s t-test and considered significant with a p-value of <0.05. The GO and KEGG pathways with FDR values (FDR<0.05) were considered statistically significant. The ceRNA network and PPI network were plotted using Cytoscape. The volcano Plot and the heatmap were generated using the EnhancedVolcano and Complex heatmap from the R (v1.3.1) BiocManager package.

## RESULTS

### CircRNA sequencing reveals the expression of thousands of circRNAs in βTC6 cells

The βTC6 cells were treated with low and high glucose for 24 h followed by total RNA isolation and RNA sequencing to identify glucose-regulated circRNAs in pancreatic β-cells (**Figure 1A**). Analysis of the RNA-seq data using CIRCexplorer2 identified more than 8000 circRNAs, among which 4,085 circRNAs were specific to low glucose (LG) and 3,015 circRNAs were specific to high glucose (HG) dataset, whereas 1,784 were common in both LG and HG. (**Figure 1B**). Analysis of circRNA distribution across different chromosomes revealed that circRNAs are generated evenly from different chromosomes except the Y chromosome (**Figure 1C**). Importantly, more than 90% of the circRNAs expressed in βTC6 cells were generated from exonic sequences termed circRNA, while a small fraction is generated from intronic sequences termed circular intronic RNAs (ciRNAs) (**Figure 1D**). Interestingly, most circRNA-generating genes in βTC6 cells generated a single circRNA, while some genes gave rise to multiple circRNAs (**Figure 1E**). Analysis of the number of exons in the exonic circRNAs revealed that most circRNAs harbor less than ten exons (**Figure 1F**).

**Figure 1:**
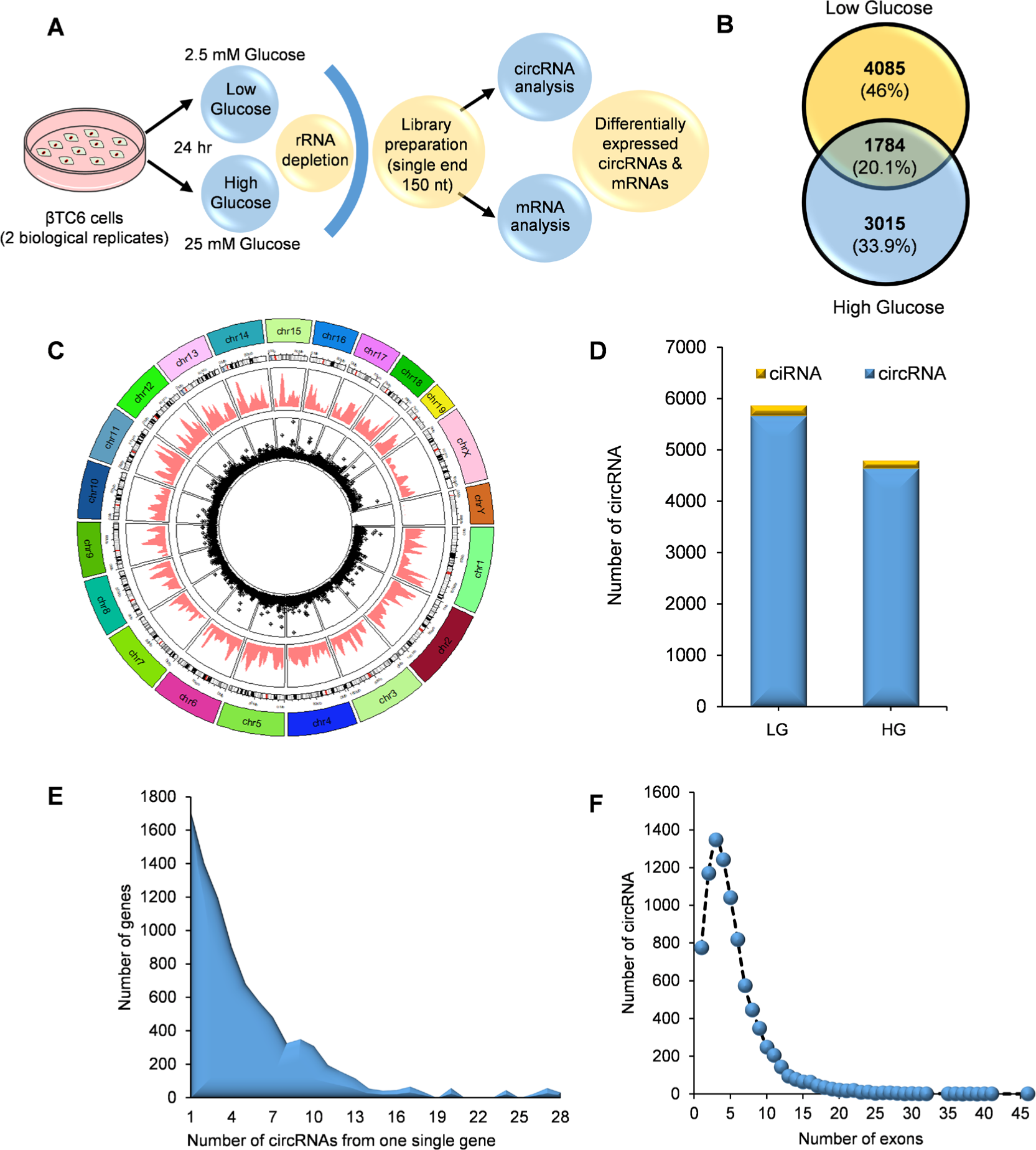
RNA-seq analysis of glucose treated βTC6 cells. **A.** The experimental design and circRNA sequencing of glucose-treated βTC6 cells. **B.** Venn diagram of circRNAs identified in LG and HG-treated βTC6 cells using CIRCexplorer2. **C.** Distribution pattern of circRNAs expressed in βTC6 cells across the chromosome. **D.** Number of exonic circRNAs and intronic ciRNAs expressed in LG and HG-treated βTC6 cells. **E.** Number of genes corresponding to the number of circRNAs generated from them. **F**. Distribution of the number of exons in exonic circRNAs in βTC6 cells.

### Validation of differentially expressed circRNAs in βTC6 cells in response to glucose

Differential expression analysis of the circRNA using the TPM method identified 325 differentially expressed circRNAs in HG compared to LG-treated βTC6 cells (**Supplementary Table S2**). To further validate the differentially expressed circRNAs, we selected six abundant and differentially expressed circRNAs, of which *circHipk2, circAW554918* are upregulated while *circHnrnpu, circCabin1, circArhgap12,* and *circRabep1* were downregulated in the HG compared to the LG treated βTC6 cells (**Figure 2A, B**). The PCR amplification of the circRNAs using divergent primers specifically amplified the target circRNAs in βTC6 cells (**Figure 2C, D**). Furthermore, Sanger sequencing of the PCR products confirmed the amplification of the backsplice junction sequence of the target circRNAs (**Figure 2E**). To confirm the circular nature of the selected circRNAs in βTC6 cells, we performed the RNase R exonuclease treatment on the total RNA from βTC6 cells and mice pancreatic islets, followed by RT-qPCR analysis of linear and circular RNA. As expected, the linear *Actb* mRNA was degraded while all the selected circRNAs were resistant to RNase R exonuclease treatment, confirming that they are true circular RNAs expressed in βTC6 cells and mice pancreatic islets (**Figure 2F, G**).

**Figure 2:**
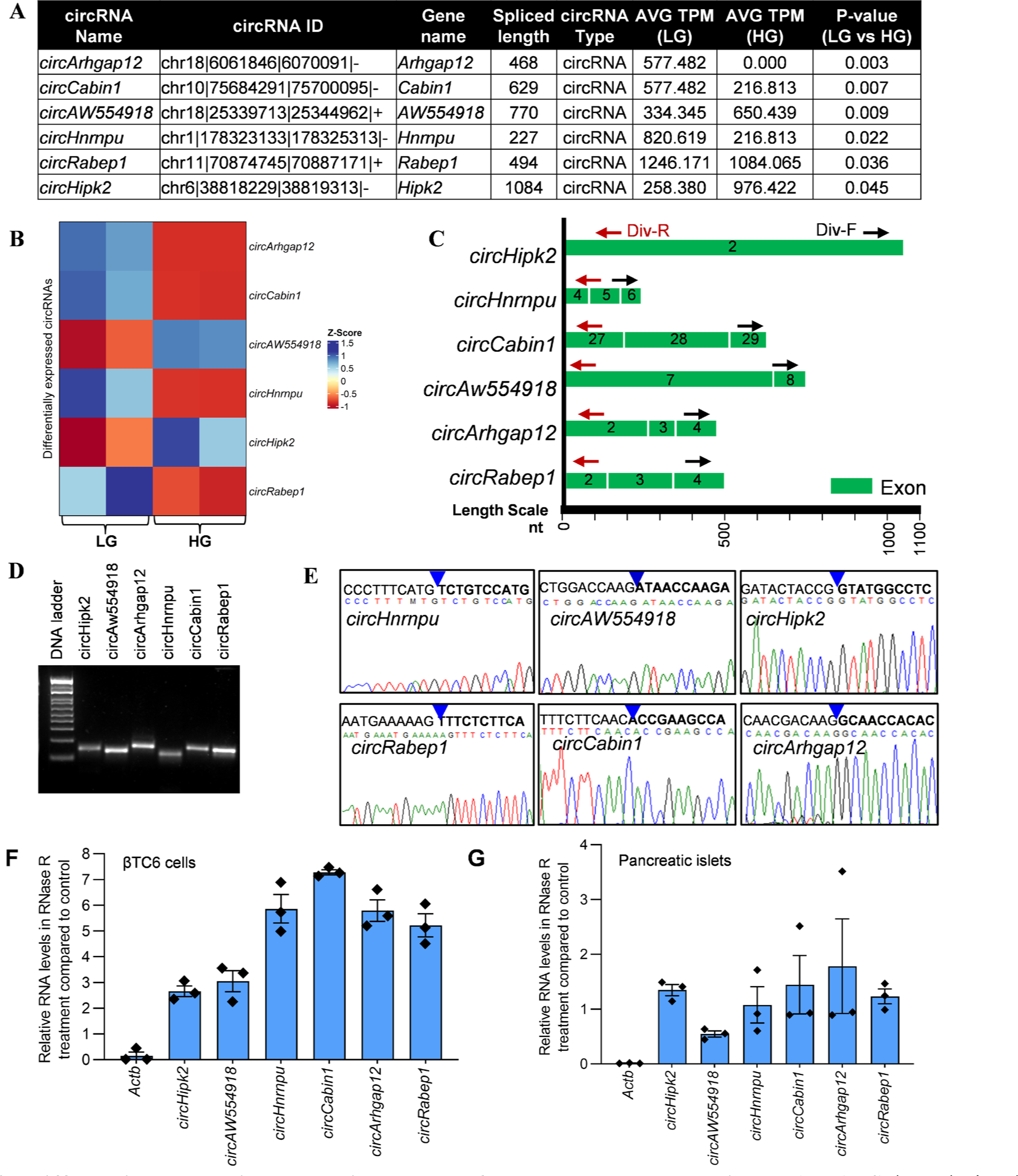
Differential expression analysis to check for glucose-regulated circRNAs. **A.** Selected circular RNA transcripts based on their significance and abundance for further analysis. **B.** Heatmap of differentially expressed circRNAs in LG and HG treated βTC6 cells. **C.** Designing of divergent primers for amplification of selected circRNAs. The arrow marks indicate the divergent primers. **D**. RT-PCR products of circRNAs amplified with divergent primers using βTC6 cells cDNA and resolved on an agarose gel stained with SYBR-Gold dye. **E**. Sanger sequencing confirms the BSJ sequence of selected circRNAs. **F-G.** RT-qPCR analysis shows linear and circRNAs levels upon RNase R treatment of total RNAs isolated from βTC6 cells (F) and mouse pancreatic islets (G). Data represent mean ± SEM of three independent repeats.

### Abundantly expressed cytoplasmic *circRabep1* is downregulated in high-glucose treated βTC6 cells

Among the differentially expressed circRNAs in βTC6 cells, we focused on *circRabep1* for further validation due to its high abundance. C*ircRabep1* is a 494 nt long exonic circRNA generated from the backsplicing of exon 2-4 of *Rabep1* pre-mRNA (NM_001291142); it is located at chr11|70874745|70887171|+ on *Rabep1* gene (**Figure 3A, Supplementary Figure S1A**). RT-qPCR expression analysis of *circRabep1* in glucose-treated βTC6 cells was consistent with RNA-seq analysis, henceforth among the differentially expressed circRNAs in βTC6 cells, we focused on *circRabep1* for further validation (**Figure 3B**). Similarly, high glucose treatment of mouse pancreatic islet for 1 day significantly reduced the expression of *circRabep1,* as seen in the case of βTC6 cells (**Figure 3C**). Consistent with the RNA-seq and RT-qPCR data, *circRabep1* copy number estimation using ddPCR suggested that βTC6 cells contain ∼150 copies in LG while HG-treated cells contain ∼100 copies per ng of RNA (**Figure 3D**). Furthermore, we determined the *circRabep1* stability by blocking the transcription by actinomycin D treatment, which led to a decrease in *Myc* mRNA levels to 10% in an hour, while the unchanged levels of *circRabep1* in βTC6 cells suggested its high stability (**Figure 3E**). Together, low expression of *circRabep1* was found to be closely associated with the high-glucose treatment of βTC6 cells. Since subcellular localization of circRNA is critical for its function, we determined the distribution of *circRabep1* in the nucleus and cytoplasm of βTC6 cells. Subcellular fractionation analysis revealed that *circRabep1* is predominantly localized in the cytoplasm of βTC6 cells suggesting that it may regulate miRNAs (**Figure 3F**).

**Figure 3.**
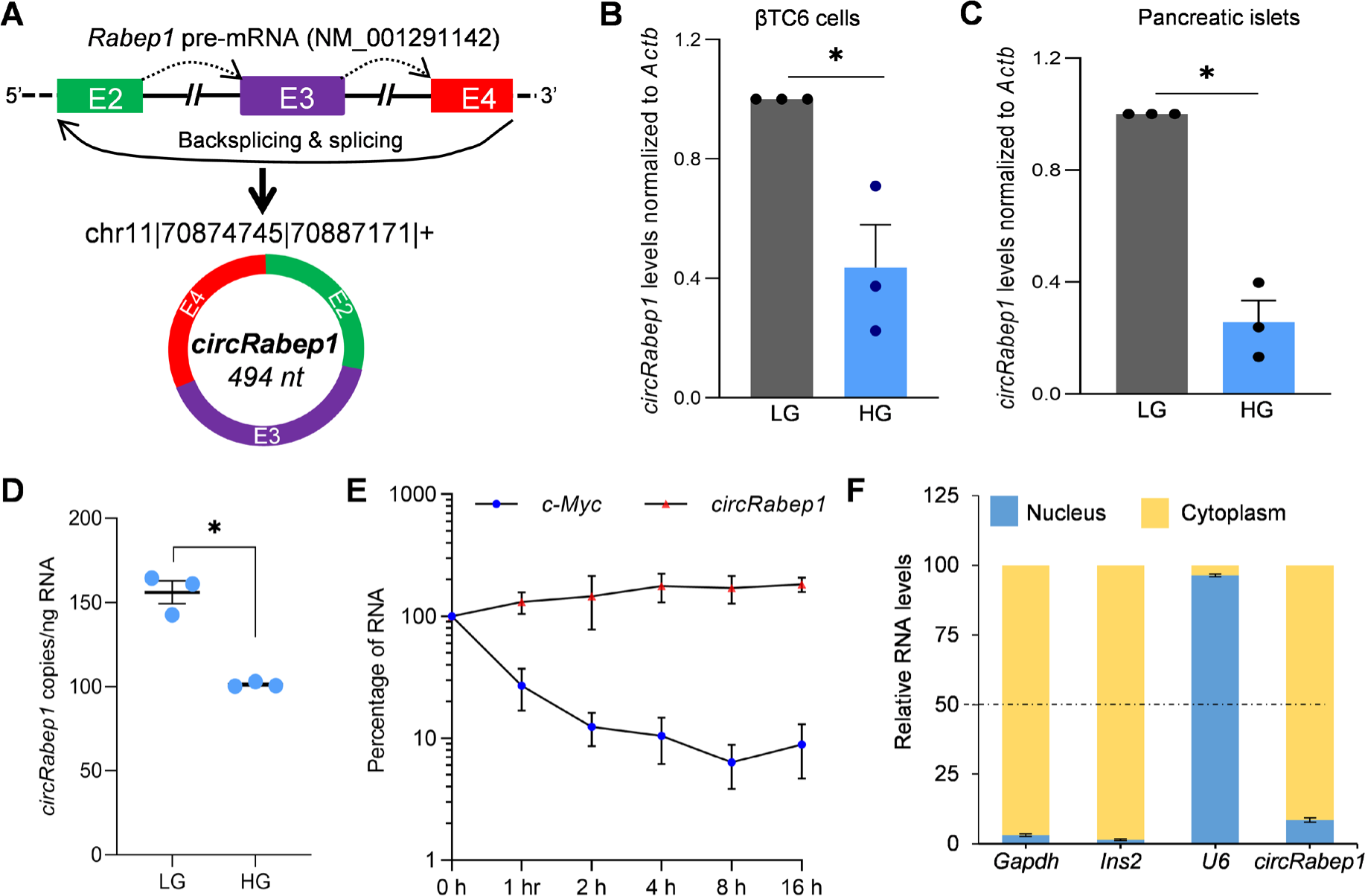
Differential expression and characterization of *circRabep1* in βTC6 cells. **A.** Schematic representation of biogenesis of *circRabep1* from the *Rabep1* pre-mRNA. **B-C.** RT-qPCR analysis of the expression levels of *circRabep1* in βTC6 cells (B) and mouse pancreatic islets (C) treated with LG or HG for 1 day. **D.** The copy number of *circRabep1* per ng of total RNA is isolated from glucose-treated βTC6 cells. **E.** RT-qPCR analysis of the stability of *Myc* mRNA and *circRabep1* in βTC6 cells by culturing the cells with actinomycin D for 16 h. **F.** Relative distribution of *circRabep1* in the nucleus and cytoplasm of βTC6 cells. The *Gapdh* mRNA and U6 snRNA serve as cytoplasmic and nuclear markers confirming the quality of subcellular fractionation. The error bars in B-F represent the means ± SEM from 3 independent experiments, and * indicates p-value <0.05.

### The *circRabep1* may regulate β-cell physiology through the circRNA-miRNA-mRNA axis

Since circRNAs regulate gene expression by acting as sponges for miRNAs, we predicted the miRNAs associated with *circRabep1*. The miRNAs were predicted to have one to three binding sites on *circRabep1* using miRDB and circAtlas ^24, 25^ (**Supplementary Figure S1B,C**). We analyzed the miRNAs targeting circRabep1 and experimentally validated miRNAs in miRTarBase to find the functional miRNAs ^26^. As shown in **Figure 4A**, analysis of the common miRNAs reported in circAtlas and miRDB, and experimentally validated mRNA targets in miRTarBase revealed 2 *circRabep1*-associated miRNAs. Importantly, two *circRabep1-*associated miRNAs, mmu-miR-125a-3p and mmu-miR-335-3p, did not alter significantly upon glucose treatment of βTC6 cells (**Figure 4B**). Furthermore, analysis of mRNA targets of these 2 miRNAs identified 155 experimentally validated targets in miRTarBase (**Supplementary Table S3, Figure 4C**).

**Figure 4:**
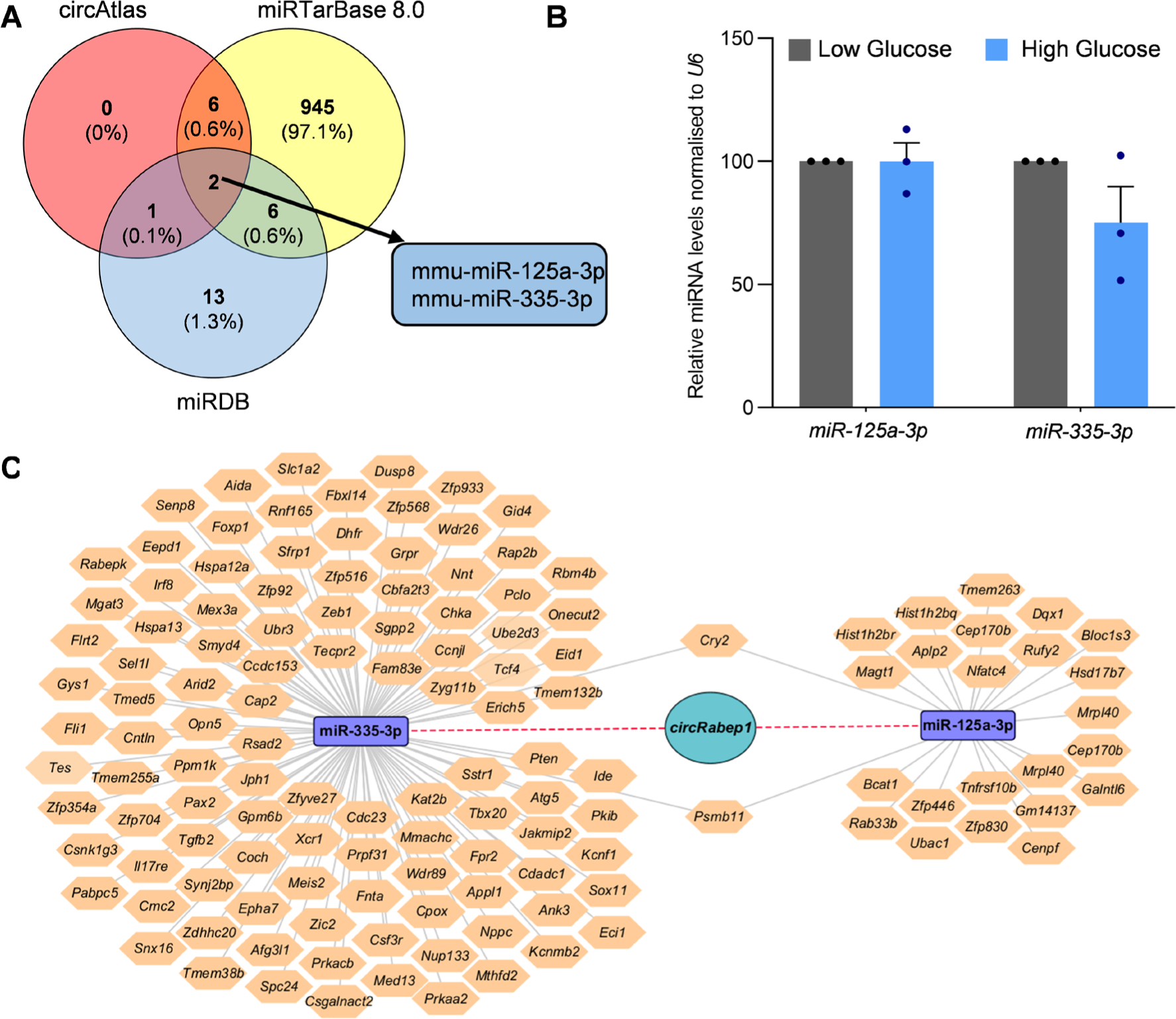
Prediction of *circRabep1*-miRNA-mRNA regulatory axis in βTC6 cells. **A.** Venn diagram of common miRNAs predicted to target *circRabep1* using miRDB, circAtlas, and miRNAs with experimentally validated mRNA targets reported in miRTarBase. **B.** RT-qPCR analysis of miRNAs targeted by *circRabep1* in LG and HG treated βTC6 cells. The error bars represent the means ± SEM from 3 independent experiments **C.** The circRNA-miRNA-mRNA regulatory network was constructed and visualized on Cytoscape. The circle node represent *circRabep1*, square nodes represent miRNAs, and hexagon nodes represent genes.

Furthermore, to find the functional significance of *circRabep1*/miRNA interaction, we predicted the Gene Ontology (GO) terms and KEGG pathways associated with the 135 genes targeted by the *circRabep1-*associated miRNAs using KOBAS webserver ^28–30^. The gene enrichment analysis of gene targeted by *circRabep1-*associated miRNAs predicted their association with various GO terms using the KOBAS database (**Supplementary Figure S2A, Supplementary Table S4**). Furthermore, computational analysis of KEGG pathways for the mRNAs associated with the *circRabep1*-miRNA network identified many highly enriched KEGG pathways (**Supplementary Figure S2B**). Finally, we analyzed the protein-protein interaction (PPI) network of genes associated with the top ten GO terms and KEGG pathways to pinpoint the regulated genes and pathways further. Interestingly, the GO function and KEGG analysis identified several genes in the PPI network critical for various cellular processes (**Supplementary Figure S2C**). These data suggest that *circRabep1* may regulate pancreatic β-cell physiology by regulating genes involved in various cellular pathways.

### PTEN is the downstream target of *circRabep1-*miRNA axis

CircRNAs act as sponges for miRNAs, thereby upregulating mRNA expression. Moreover, the levels of circRNAs are positively correlated with the downstream mRNA targets of the circRNA-associated miRNAs through the ceRNA regulatory network. To identify the downstream target gene of *circRabep1* in pancreatic β-cell, we identified the mRNAs differentially expressed upon treatment of βTC6 cells with high glucose. As shown in **Figure 5A**, 557 genes were differentially expressed in βTC6 cells treated with high glucose for 24 hr compared to the low glucose treated sample (**Supplementary table S5**). To identify the potential genes regulated by *circRabep1* in glucose-treated βTC6 cells, we analyzed differentially expressed genes and mRNA targets of *circRabep1-*associated miRNAs. Interestingly, four genes, including *Tes, Tcf4, Ubed3,* and *Pten* were downregulated in high glucose-treated βTC6 cells and are targets of *circRabep1-*associated miRNAs (**Figure 5A**). Although *circRabep1* downregulation in the HG-treated βTC6 cells positively correlated with the decrease in the four downstream targets in RNA-seq, RT-qPCR analysis of mRNAs only validated the positive correlation with *Pten* mRNA expression, and *Tcf4* was undetected (**Figure 5B**). Moreover, the *Pten* mRNA has been predicted to be targeted by *circRabep1-*associated miRNA, miR-335-3p (**Figure 4C**). As *circRabep1-*miR-335-3p association was predicted by the miRDB and circAtlas web tools, we wanted to validate the direct interaction of *circRabep1* and miR-335-3p by pulling down endogenous *circRabep1* using biotinylated-ASO targeting the *circRabep1* BSJ sequence (**Figure 5C**). As shown in **Figure 5D**, *circRabep1* was significantly enriched in *circRabep1* pulldown samples compared to the control biotinylated-ASO pulldown samples in βTC6 cells. Similarly, miR-335-3p was significantly enriched in the *circRabep1* pulldown samples supporting the existence of *circRabep1*–miR-335-3p complexes in βTC6 cells (**Figure 5E**).

**Figure 5:**
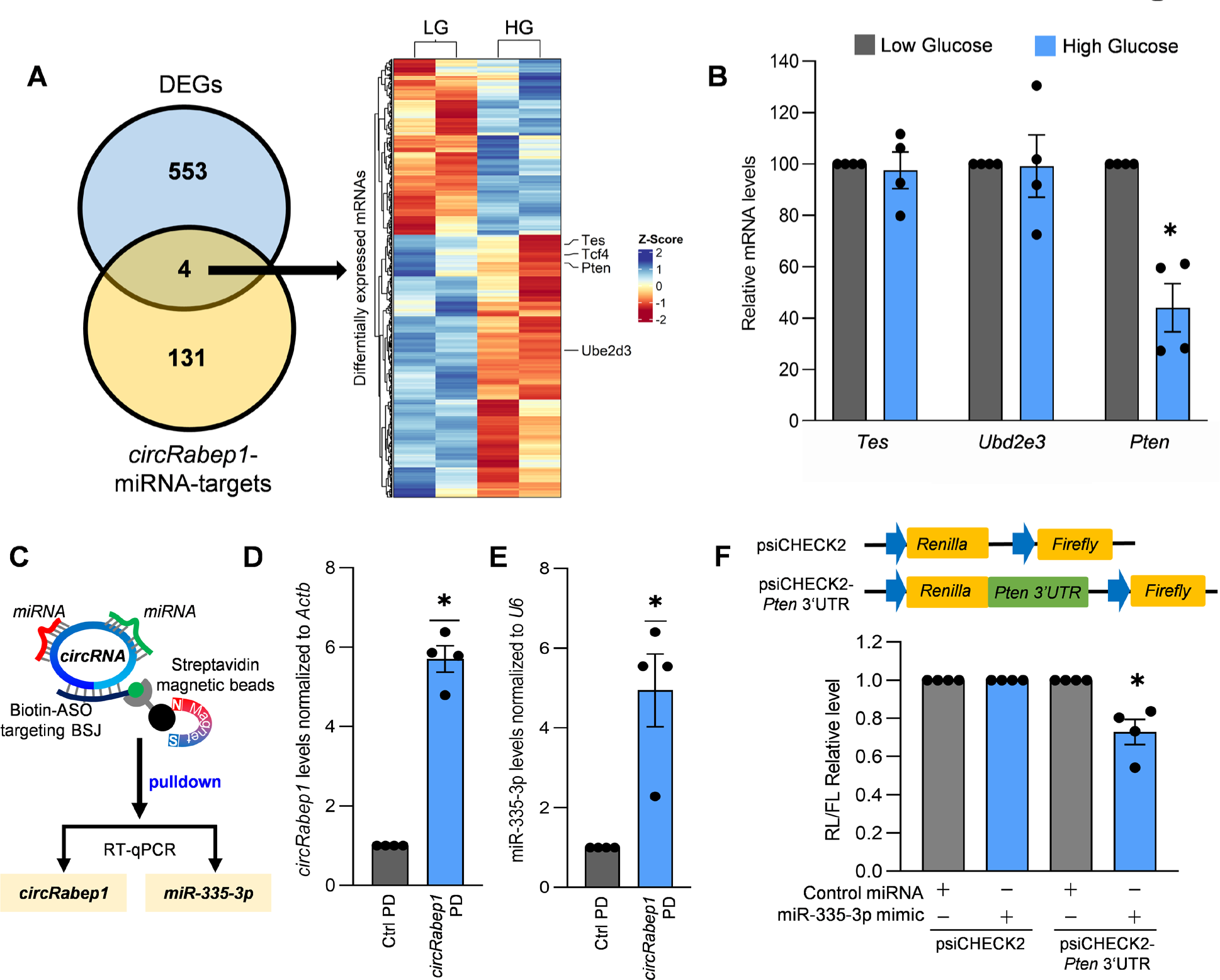
*CircRabep1* regulates PTEN expression by binding to miR-335-3p. **A.** Venn diagram depiction of the common genes between the *circRabep1* targets and DE-mRNAs (*left*). Heatmap showing the 5 genes differentially expressed upon glucose treatment and are targets of *circRabep1*-miRNA axis (*Right*). **B.** RT-qPCR analysis of mRNA targets of miR-335-3p in LG and HG treated βTC6 cells. **C.** Schematic representation of *circRabep1* pulldown using biotin-labeled ASO targeting the BSJ sequence of *circRabep1.* **D-E.** RT-qPCR and enrichment analysis of *circRabep1* (D) and miR-335-3p (E) in the pulldown samples using biotin-labeled ASO targeting *circRabep1* BSJ sequence compared to control biotin-labeled ASO. **F.** Top, schematic of psiCHECK2 reporter expressing renilla luciferase (RL) and internal control firefly luciferase (FL); the psiCHECK2-Pten 3’UTR vector contains a fragment of *Pten* 3’UTR sequence targeted by miR-335-3p downstream of the renilla luciferase sequence. Bottom, the relative ratio of RL/FL activity of βTC6 cells transfected with either miR-335-3p mimic or control miRNA and either psiCHECK2 or psiCHECK2-*Pten* 3’UTR were plotted. Data in B, D-F represent the means ± SEM from 4 independent experiments, and * indicates p-value <0.05.

As shown in **Supplementary Figure S3,** miR-335-3p has 3 target sites on the 3’ UTR of *Pten* mRNA predicted by miRTarBase web tool using MiRanda, TargetScan, and miRDB ^24, 26, 27^. Since the first site was having higher score and was predicted by all three databases, we cloned that sequence in the psiCHECK2 vector downstream to renilla. Furthermore, co-transfection of miR-335-3p mimic along with psiCHECK2-Pten 3’ UTR vector reduced the luciferase expression normalized to the control psiCHECK2 plasmid. The specific inhibition of luciferase activity by miR-335-3p indicates direct interaction of *Pten* 3’UTR with miR-335-3p, which may inhibit PTEN expression in βTC6 cells (**Figure 5F, Supplementary Figure S3**). Since *circRabep1* is associated with miR-335-3p which targets *Pten* 3’UTR, we concluded that *circRabep1* may promote PTEN expression by inhibiting miR-335-3p activity in βTC6 cells.

### The *circRabep1* promotes PTEN expression in pancreatic β-cells

Importantly, silencing of *circRabep1* in βTC6 cells decreased *Pten* mRNA and PTEN protein levels, suggesting the regulation of PTEN through inhibiting *Pten*-associated miRNAs by *circRabep1* (**Figure 6A, B**). To determine the molecular mechanism through which *circRabep1* regulates *Pten* mRNA expression, we analyzed the stability and translation efficiency of *Pten* mRNA expression upon *circRabep1* silencing in βTC6 cells. Since *circRabep1* silencing decreased *Pten* mRNA expression, we hypothesized that *circRabep1* might regulate *Pten* mRNA stability. We determined the differences in *Pten* mRNA stability by comparing the half-lives of *Pten* mRNA by blocking the transcription using actinomycin D with or without *circRabep1* silencing in βTC6 cells. As shown in **Figure 6C**, the half-life of *Gapdh* mRNA was not altered upon *circRabep1* silencing. However, there was a specific decrease in the half-life of *Pten* mRNA to 1 h in *circRabep1* silenced cells compared to 1.8 h in control cells. These data suggest that the decrease in *Pten* mRNA stability could be due to the higher availability of miR-335-3p in the *circRabep1* silenced condition (**Figure 6C**).

**Figure 6.**
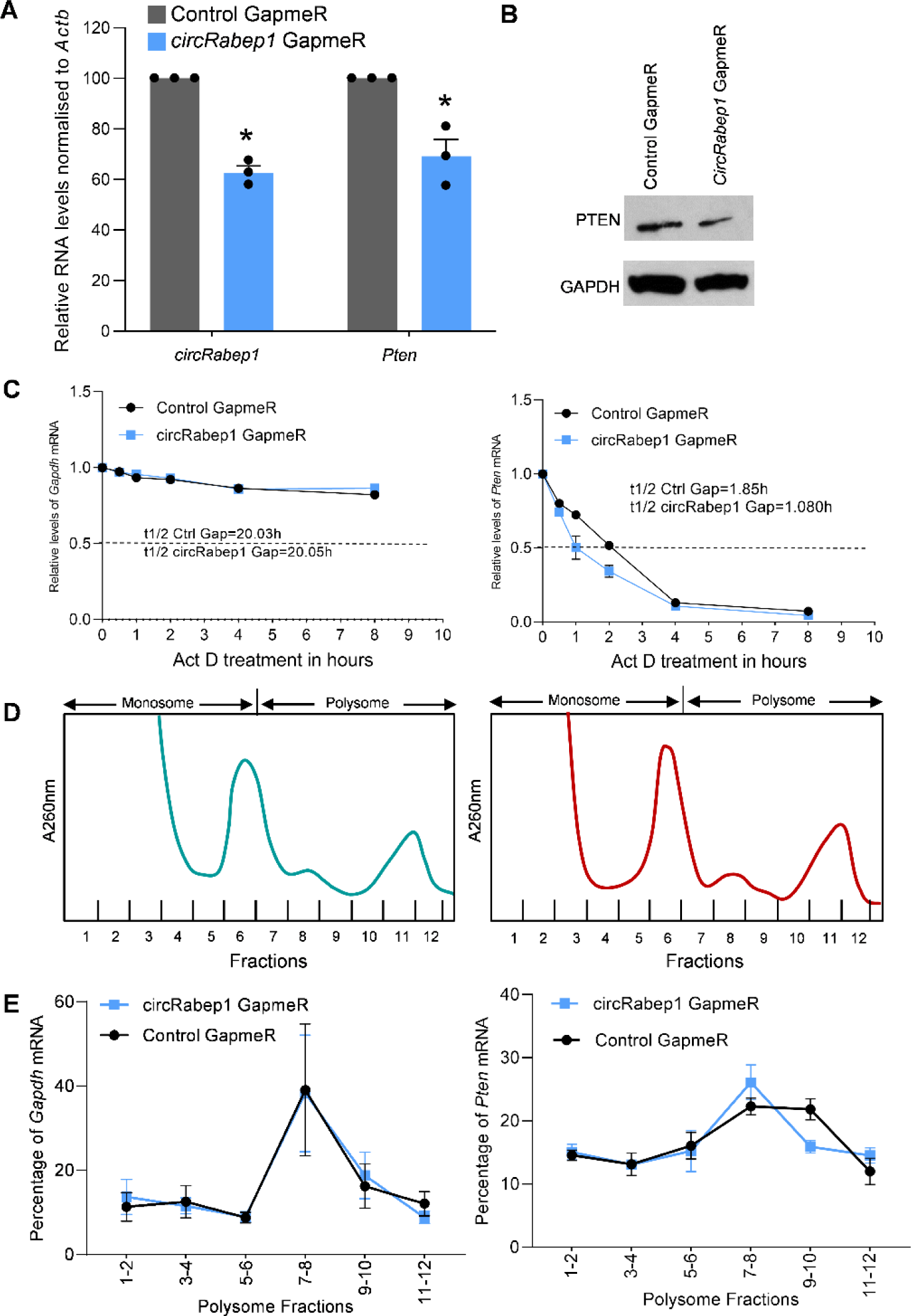
Regulation of *Pten* mRNA stability and translation by c*ircRabep1.* **A.** RT-qPCR analysis of *Pten* mRNA expression in βTC6 cells upon *circRabep1* knockdown. **B.** Western blot analysis of PTEN expression in βTC6 cells upon *circRabep1* knockdown. **C.** The *Pten* and *Gapdh* mRNA stability were assessed by treating the βTC6 cells with actinomycin D to block transcription with or without *circRabep1* silencing. RT-qPCR analysis was used to find the half-lives of these mRNAs. **D-E.** Polyribosomes in cytoplasmic extracts of βTC6 cells were fractionated through sucrose gradients (D), and the relative distribution of *Pten* and housekeeping *Gapdh* mRNA in different fractions was measured by RT-qPCR analysis and represented as the percentage of total RNA in the gradient (E). The error bars represent the mean ± SEM of three independent experiments.

Since miRNAs can regulate mRNA translation along with stability, we sought to analyze the translation efficiency of *Pten* mRNA in βTC6 cells upon *circRabep1* silencing. Representative sucrose gradient polyribosome profiles indicated no significant changes in global translation upon *circRabep1* silencing (**Figure 6D**). The sucrose gradient fractions 1-6 were considered as monosome or non-translating pool, while fractions 7-12 contained actively translating polyribosomes or polysomes. We analyzed the distribution pattern of *Pten* mRNA and housekeeping *Gapdh* mRNA in the polyribosome fractions with or without *circRabep1* silencing. As shown in **Figure 6E**, *Pten* mRNA levels were low in the non-translating monosome fractions and abundant in the translating polysome fractions, indicating that both *Gapdh* and *Pten* mRNAs were actively translated in βTC6 cells. Importantly, the silencing of *circRabep1* shifted the *Pten* mRNA towards lighter polysomes while the polysome association of housekeeping *Gapdh* mRNA was unchanged, suggesting a specific inhibition of *Pten* mRNA translation by higher bioavailability of miR-335-3p upon *circRabep1* silencing (**Figure 6E**). Together, these data support the hypothesis that miR-335-3p binds to *Pten* mRNA and suppresses the stability and translation of *Pten* mRNA, while *circRabep1* competes with *Pten* mRNA for binding to miR-335-3p, promoting PTEN expression.

### The *circRabep1* promotes β-cells growth by regulating PTEN in high glucose condition

Glucose treatment of β-cells has been known to induce β-cell growth and proliferation. Since *Pten* is a well-known cell growth and proliferation regulator, we asked if *circRabep1* regulates β-cell growth by modulating miR-335-3p activity. Importantly, inhibition of miR-335-3p in βTC6 cells using the miR-335-3p inhibitor resulted in a significant increase in *Pten* mRNA, suggesting the regulation of PTEN by miR-335-3p (**Figure 7A**). Furthermore, western blot analysis revealed that miR-335-3p inhibition using miR-335-3p antagomir led to an upregulation of PTEN levels in βTC6 cells (**Figure 7B**). As shown in **Figure 7C**, inhibition of miR-335-3p, which increases PTEN expression, led to a significant decrease in cell viability and proliferation of βTC6 cells measured by WST-1 assay. Furthermore, *circRabep1* silencing in βTC6 cells increases the availability of miR-335-3p and suppresses PTEN expression, promoting cell survival and proliferation. Importantly, the proliferative phenotype seen in *circRabep1* silencing was rescued by inhibiting miR-335-3p in *circRabep1* silenced cells (**Figure 7C**). Since the steady-state levels of miR-335-3p do not change significantly in glucose-treated βTC6 cells and *circRabep1* levels positively correlate with PTEN expression, we propose that *circRabep1* may regulate pancreatic β-cell growth at least in part by interacting with the miR-335-3p (**Figure 7D**).

**Figure 7.**
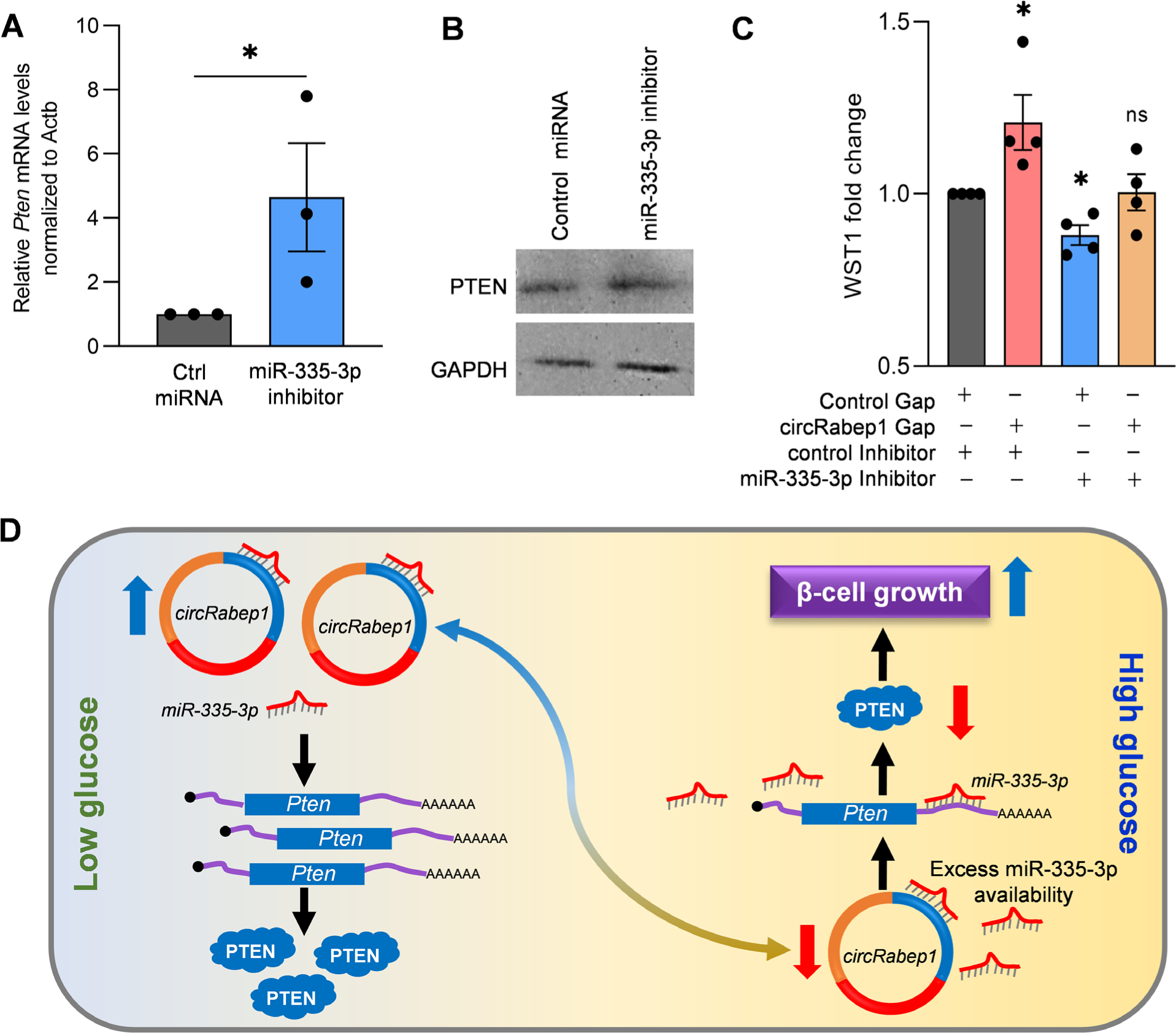
Regulation of β-cell growth by *circRabep1*-miR-335-3p-PTEN axis. **A.** RT-qPCR analysis of *Pten* mRNA expression in βTC6 cells upon miR-335-3p inhibition. **B.** Western blot analysis of PTEN in βTC6 cells transfected with control and miR-335-3p antgomir. **C.** WST-1 assay evaluating the βTC6 cells growth and proliferation in *circRabep1* silencing condition with or without miR-335-3p inhibitor. **D.** Proposed model whereby *circRabep1* acts as a ceRNA for miR-335-3p; *circRabep1* promotes the expression of PTEN by sponging miR-335-3p in low glucose condition inhibiting cell proliferation, while decreased *circRabep1* expression in high glucose condition increased the availability of functional miR-335-3p suppressing PTEN expression, which in turn promotes β-cell growth. The error bars in A and C represent the mean ± SEM of 3-4 independent experiments.

## DISCUSSION

Despite the lack of protein-coding ability, circRNAs have emerged as an important regulator of gene expression ^12, 31, 32^. CircRNAs are involved in cell proliferation, cell development, differentiation, and cell survival by modulating gene transcription, mRNA stability, translation, and protein function ^12, 32, 33^. CircRNAs are also associated with various diseases, including autoimmune disorders, Alzheimer’s, cancer, and diabetes ^33, 34^. The recent discovery of thousands of circRNAs in pancreatic islets and their dysregulation in conditions associated with T2D indicates the possibility that they are key regulators of pancreatic β-cell physiology ^8, 9^. However, only a few recent studies investigated the possible role of a few circRNAs in β-cell function and the development of diabetes ^9, 16, 35^. Since we are starting to understand the role of circRNAs in pancreatic β-cell physiology, more studies are required for a better understanding of these novel transcripts in β-cells during the development of T2D.

Dysfunction of pancreatic β-cells and impaired insulin secretion in response to glucose leads to elevated blood glucose levels and T2D ^3^. The β-cell response to external glucose is tightly regulated by various genes regulated at transcriptional and posttranscriptional levels ^36^. Long-term palmitate and glucose exposure have been shown to alter β-cell gene expression and impair normal β-cell function ^3, 4^. However, the glucose-regulated circRNA transcriptome and its function in β-cell have not been studied. In this study, we used RNA-seq technology to identify circRNAs expressed in βTC6 cells treated with glucose. We identified more than 8000 circRNAs expressed in the βTC6 cells, and most of the circRNAs were generated from the exonic sequences (**Figure 1**). Interestingly, differential expression analysis and molecular validation experiments identified several differentially expressed circRNAs, including *circRabep1.* Since *circRabep1* was predominantly localized in the cytoplasm, it could regulate cellular functions by translating into polypeptides or interacting with RBPs or miRNAs. However, *circRabep1 lacks* the protein-coding ability, as reported by the riboCIRC database ^37^. In addition, analysis of RBP binding sites on *circRabep1* using RBPmap web server revealed numerous binding sites for many RBPs (data not shown) ^38^. Since selecting the right RBP for regulation was challenging, we focused on the *circRabep1*-miRNA regulatory axis. As shown **Figure 3**, we could predict two miRNAs with functional mRNA targets, including miR-335-3p. As the ceRNA hypothesis suggests a positive correlation between the circRNA and the downstream mRNA expression through inhibition of miRNA, we sought to identify common genes that are downregulated in high glucose-treated βTC6 cells and are targets of *circRabep1.* We could identify a few genes, including PTEN, as a downstream target of the *circRabep1-*miR-335-3p axis (**Figure 5**). However, the changes in the expression of *Pten* and other mRNAs in RNA-seq were moderate, which could be due to moderate but significant downregulation of *circRabep1* in high glucose-treated βTC6. Importantly, *circRabep1-*silencing suppressed PTEN expression significantly in βTC6 cells by suppressing *Pten* mRNA translation and stability (**Figure 6**). Moreover, *circRabep1* silencing promoted βTC6 cell growth, while miR-335-3p inhibition rescued the *circRabep1* effect on cell growth (**Figure 7**). Our data identified the novel *circRabep1-*miR-335-3p-PTEN regulatory axis that regulates pancreatic β-cell growth.

PTEN was first discovered as a tumor suppressor with loss of expression in various cancers ^39^. Additional studies established that PTEN inhibits the well-known PI3K/AKT pathway for cell growth and survival ^40, 41^. PTEN is a lipid phosphatase that converts PIP3 to PIP2 by removing the 3’ phosphate. PI3K adds the 3’ phosphate to convert PIP2 to PIP3. PI3K is activated in response to growth factors leading to the accumulation of PIP3, which activates kinases with PH domains like AKT to induce cell growth and proliferation ^40^. Since PTEN antagonizes PI3K-mediated cell growth, loss of PTEN in cancers or pancreatic β-cells has been shown to promote cell growth and survival ^41, 42^. Importantly, several reports established that PTEN inhibits β-cell function, growth, and survival ^42, 43^. Although deletion or downregulation of PTEN often leads to tumor development, reduced expression of PTEN in β-cells leads to activation of AKT, promoting β-cell regeneration and resistance to streptozotocin-induced diabetes in mice ^43^. The downregulation of PTEN in high glucose treated β-cells by *circRabep1*-miR335-3p axis is a novel mechanism for regulation of β-cell growth and survival in response to short-term high glucose. Although our study identified a novel mechanism of *circRabep1* mediated β-cell growth, *in vivo* experiments in animal models would provide useful data for therapeutic use in diabetes. It is important to note that we predicted the *circRabep1-*associated miRNAs using miRDB and downstream targets through miRTarBase. Other programs could predict different sets of miRNAs and different downstream targets. Since *circRabep1* have many downstream targets in the miRNA axis, the effect of *circRabep1* on pancreatic β-cell growth could be partly through other mechanisms, which needs further investigation.

In summary, we identified glucose-regulated circRNAs in the pancreatic β-cells for the first time. This study identified *circRabep1* as a downregulated circRNA in the high glucose treated βTC6 cells and identified the *circRabep1*-miR-335-3p-PTEN regulatory network. These findings may serve as a valuable resource for further investigation of circRNAs in pancreatic β-cell physiology. Further studies with extensive circRNA profiling and uncovering the downstream regulatory network in pancreatic β-cell will help develop novel therapies that can potentially restore β-cell function in diabetes.

## DECLARATIONS

### Data availability

All the data generated in this study are included in the main text or supplementary data. The RNA-seq data generated in this study were deposited in the European Nucleotide Archive (ENA) with Accession numbers PRJEB64535.

### Author contributions

Debojyoti Das: Conceptualization, Methodology, Investigation, Formal analysis, Validation, Visualization, Writing—original draft. Sharmishtha Shyamal: Methodology, Investigation, Formal analysis, Data Curation, Validation, Visualization, Writing—original draft. Smruti Sambhav Mishra: Investigation, Formal analysis, Validation. Susovan Sadhukhan: Investigation, Formal analysis, Writing— review & editing. Amaresh C. Panda: Conceptualization, Methodology, funding acquisition, Supervision, Writing— review & editing

## Acknowledgements

We thank our lab colleagues for the helpful discussion and for proofreading the manuscript.

## Ethics approval and consent to participate

Animal studies were conducted with the approval of the Institutional Animal Ethics Committee of Institute of Life Sciences, Bhubaneswar, India.

## Consent for publication

All authors consent this manuscript for publication.

