## Supplementary Figures for "Glucose-regulated circular RNA Rabep1 regulates pancreatic β-cell growth by modulating miR-335-3p/PTEN axis"

**Supplementary Tables and Figures**

**Supplementary Table S1:** List of oligonucleotides used for the study

**Supplementary Table S2:** circRNA annotation in ßTC6 cells using CIRCexplorer2.

**Supplementary Table S3:** Target genes of *circRabep1*-associated miRNAs

**Supplementary Table S4:** GO and KEGG analysis of genes targeted by *circRabep1*-miRNA axis

**Supplementary Table S5:** Differentially expressed mRNAs in glucose treated ßTC6 cells

### Das et al Fig. S1

**A** ***circRabep1* sequence**

TTTCTCTTCAGCAACGGGTAGCAGAATTGGAAAAAATTAATGCAGAATTTTTACGTGCACAGCAG
CAGCTTGAGCAAGAATTTAATCAAAAGAGAGCAAAATTTAAGGAGCTCTATTTGGCTAAAGAGGA
GGATCTGAAGAGGCAAAATGCAGTATTGCAAGCTGCACAGGACGATTTGGGTCACCTTCGGACA
CAGCTGTGGGAGGCTCAGGCAGAGATGGAGAACATTAAGGCAATTGCCACAGTCTCTGAGAATA
CTAAGCAAGAAGCTATAGATGAAGTTAAAAGGCAATGGAGAGAAGAAGTTGCTTCACTTCAGGCT
ATAATGAAAGAGACAGTCCGTGACTATGAGCATCAGTTTCACCTGAGGCTGGAGCAGGAGCGAG
CACAGTGGGCACAGTATCGAGAATCTGCAGAGCGGGAAATAGCTGACTTAAGAAGAAGGTTGTC
TGAAGGTCAAGAGGAAGAAAATTTAGAAAATGAAATGAAAAAG

**B**

**Targets of *circRabep1* predicted by miRDB**

| miRNA Name | No. of Target sites | Seed Location | Target Score |
|---|---|---|---|
| miR-7063-3p | 1 | 91 | 94 |
| miR-6973b-3p | 1 | 91 | 94 |
| miR-6400 | 2 | 73, 262 | 89 |
| miR-7014-3p | 2 | 89, 329 | 83 |
| miR-130b-5p | 2 | 89, 329 | 83 |
| miR-153-5p | 2 | 478, 484 | 83 |
| miR-20a-3p | 2 | 39, 147 | 83 |
| miR-217-5p | 1 | 148 | 70 |
| miR-335-3p | 1 | 486 | 69 |
| miR-7090-5p | 1 | 248 | 67 |
| miR-7030-3p | 1 | 93 | 67 |
| miR-511-5p | 2 | 231, 286 | 65 |
| miR-7052-3p | 1 | 212 | 61 |
| miR-7221-5p | 1 | 331 | 61 |
| miR-6961-3p | 2 | 123, 460 | 60 |
| miR-12183-5p | 3 | 300, 437, 439 | 57 |
| miR-1903 | 3 | 300, 437, 439 | 56 |
| miR-6963-3p | 1 | 138 | 54 |
| miR-544-3p | 1 | 41 | 53 |
| miR-3057-3p | 1 | 198 | 52 |
| miR-6946-3p | 3 | 299, 438, 465 | 50 |
| miR-125a-3p | 1 | 362 | 50 |

**C**

**Targets of *mmu-Rabep1_0001* predicted by circAtlas**

| microRNA name | # sites by PITA | # sites of miRanda | # sites of targetScan |
|---|---|---|---|
| mmu-miR-3059-5p | 0 | 1 | 1 |
| mmu-miR-6516-3p | 0 | 1 | 1 |
| mmu-miR-5125 | 0 | 1 | 1 |
| mmu-miR-188-3p | 0 | 1 | 1 |
| mmu-miR-125a-3p | 0 | 1 | 1 |
| mmu-miR-1904 | 0 | 1 | 2 |
| mmu-miR-329-5p | 0 | 1 | 1 |
| mmu-miR-335-3p | 0 | 1 | 1 |
| mmu-miR-6400 | 0 | 1 | 2 |

**Supplementary Figure S1: miRNA targets of *circRabep1*. A.** Spliced sequence of *circRabep1*. **B-C.** List of miRNAs targeted by *circRabep1* predicted using custom prediction tool of miRDB (B) and circAtlas (C) web server.

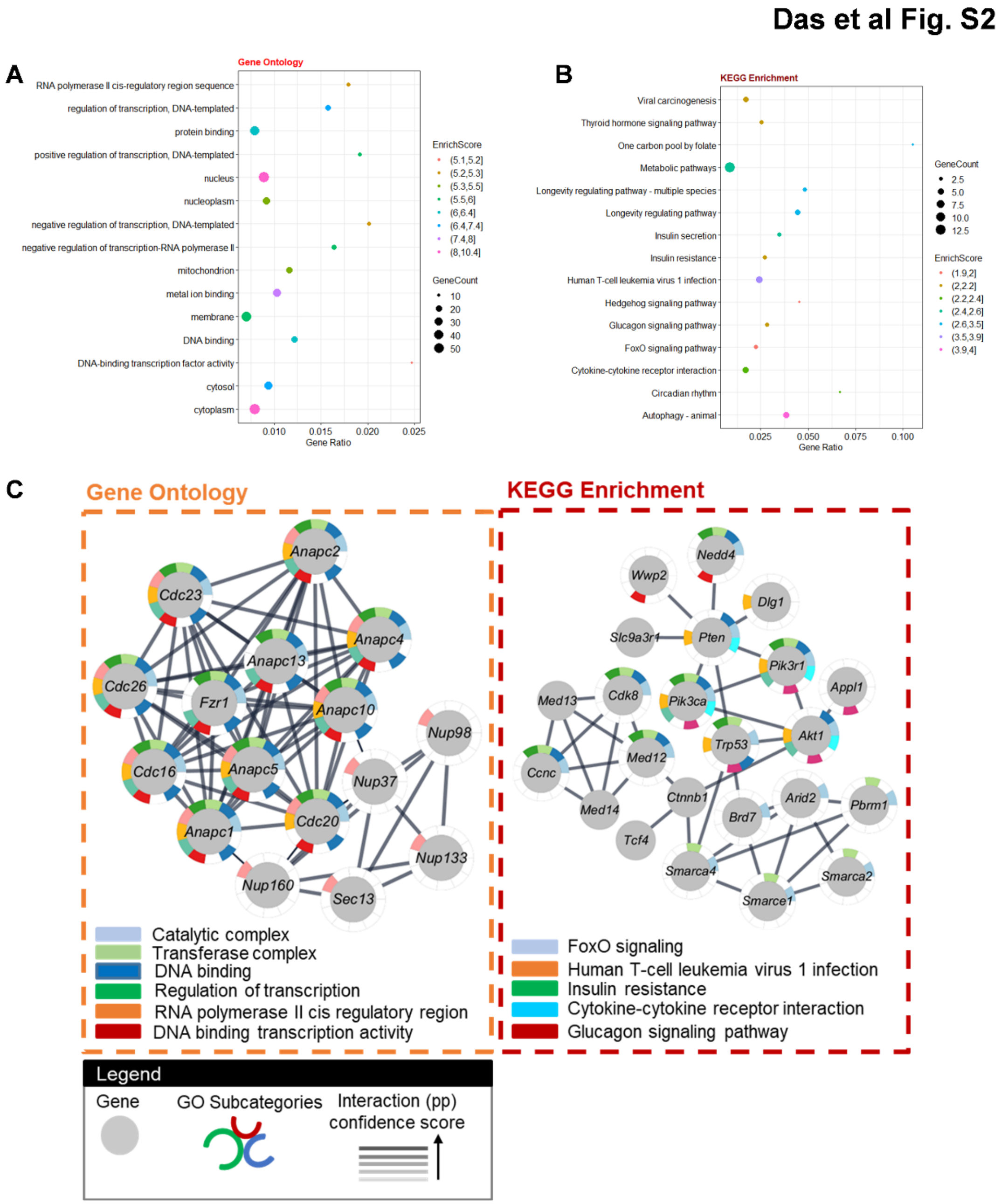


**Supplementary Figure S2. A.** Gene ontology (KOBAS 3.0) analysis of genes targeted by miRNAs associated with *circRabep1*. **B.** KEGG pathway enrichment analysis of the genes that are associated with *circRabep1*-miRNA-mRNA regulatory axis. **C.** Gene ontology and KEGG analysis of the genes in the top 15 GO terms and KEGG pathways. Gray circles represent the genes in the PPI networks; outside colored layers illustrate the different GO terms, and the gray lines represent the confidence score of the PPI network.

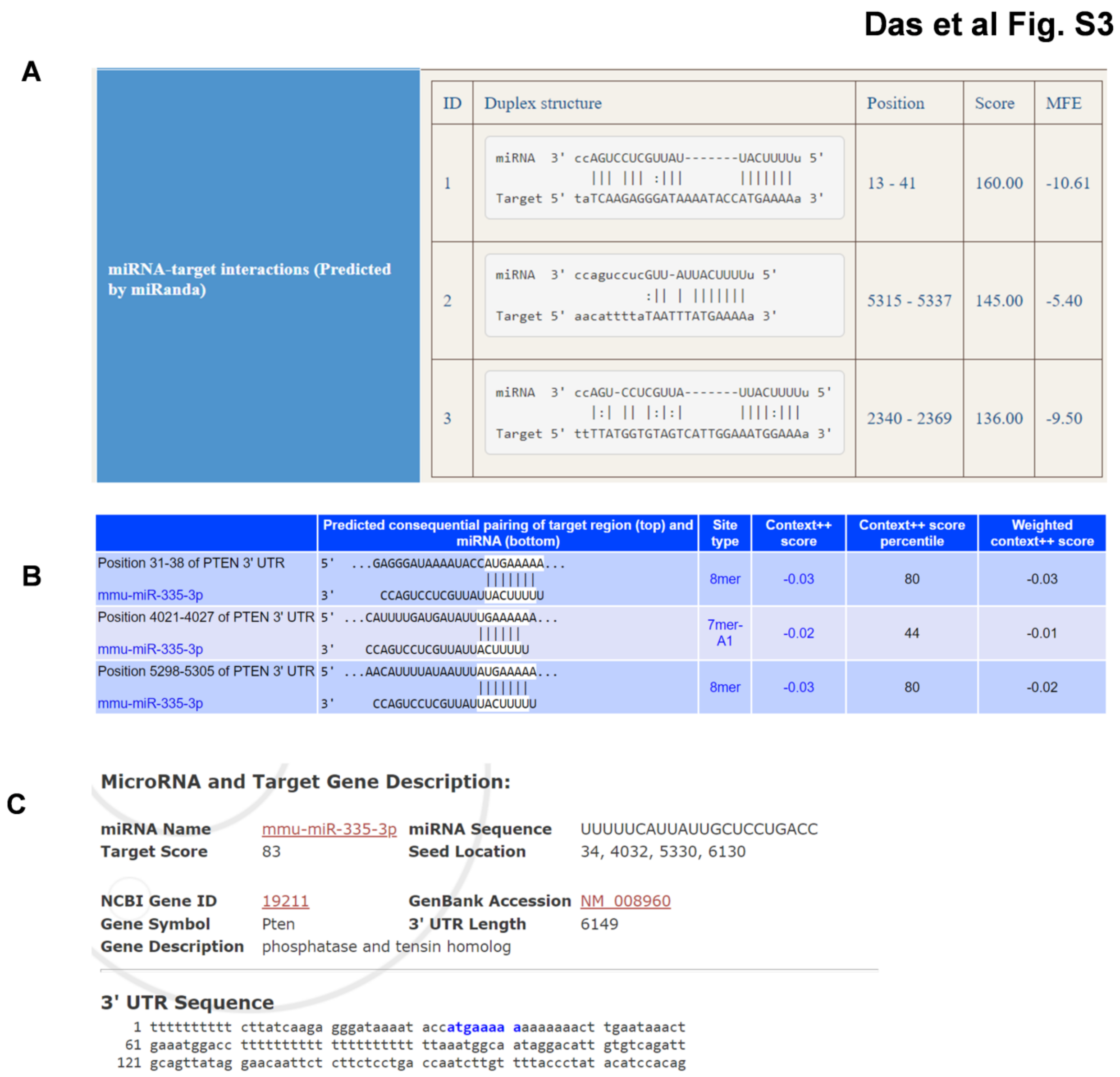


**Supplementary Figure S3. A-C.** Screenshots of the interaction of miR-335-3p with the 3′ UTR of mouse *Pten* mRNA predicted by miRTarBase using miRanda (A), TargetScan (B), and miRDB (C).
